# Cystic proliferation of embryonic germ stem cells is necessary to reproductive success and normal mating behavior in medaka

**DOI:** 10.1101/2020.08.30.274480

**Authors:** Luisa F. Arias Padilla, Diana C. Castañeda-Cortés, Ivana F. Rosa, Rafael Henrique Nóbrega, Juan I. Fernandino

**Affiliations:** Instituto Tecnológico de Chascomús, INTECH (CONICET-UNSAM), Chascomús, Argentina; Reproductive and Molecular Biology Group, Department of Structural and Functional Biology, Institute of Biosciences, São Paulo State University (UNESP), Botucatu, São Paulo, Brazil

**Keywords:** Ndrg1, Mitosis, Gametogenesis, Reproduction, CRISPR/Cas9, Mating vigor

## Abstract

The production of an adequate number of gametes in both sexes is necessary for normal reproduction, for which the regulation of proliferation from early gonadal development to adulthood is key. Cystic proliferation of embryonic stem germ cells prior the onset of gametogenesis is an especially important step prior to the beginning of meiosis. However, in vertebrates, the molecular regulators of cystic proliferation remain unknown. Here, we report that *ndrg1b*, a member of the N-myc downstream regulated family, is an important regulator of cystic proliferation in medaka. We generated mutants of *ndrg1b* that led to a disruption of proliferation type II, independently of the TGF-β signaling pathway. This loss of cystic proliferation was observed from embryogenic to adult stages, impacting the success of gamete production and reproductive parameters such as spawning and fertilization. Interestingly, the depletion of cystic proliferation of the *ndrg1b* mutant also impacted male sexual behavior, with a decrease of mating vigor. These data illustrate why it is also necessary to consider gamete production capacity in order to analyze reproductive behavior.

**HIGHLIGHTS:**

- Ndrg1b is involved in the regulation of cystic proliferation in gonad from embryo to adulthood.
- The cystic proliferation is independently of the TGF-β signaling pathway.
- Decrease of production of gametes declines reproductive success for both sexes.
- Reduction of cystic proliferation declines male sexual behavior, with a decrease of mating vigor.

## INTRODUCTION

Cell division and differentiation are well-controlled processes for the development of a functional organ in multicellular organisms. In gonads, these processes are also essential to guarantee the correct production of gametes necessary for successful reproduction (Lombardi, 1998). Vertebrate gonads acquire sex-specific features during gonadogenesis, an exceptional, complex, multistep process that includes migration of primordial germ cells (PGCs) to reach the gonadal primordium, proliferation, differentiation and meiosis (DeFalco and Capel, 2009; Richardson and Lehmann, 2010). Defects in any of these steps may lead to sex abnormalities including infertility and sex reversal.

Proliferation of embryonic germ stem cells (EGSCs, the name applied to PGCs after they reach the gonadal primordium) originates a cohort of germ cells that will ultimately form the primary population of germ cells in the gonads of both sexes (Tanaka, 2019). This type of slow, intermittent division, also called type I proliferation, is similar to the self-renewal of germline stem cells in the adult ovary or testis (Chen and Liu, 2015; Nakamura *et al*., 2010), and it is responsible for ensuring the production of the correct number of germline stem cells that will initiate gametogenesis (Tanaka, 2019). In addition, the hyperproliferation of mitotically active germ cells in *hotei* medaka, a mutant fish for anti-Müllerian hormone receptor II (*amhrII*), induces 50% male-to-female sex reversal and loss of reproductive success in adult fish (Morinaga *et al*., 2007; Nakamura *et al*., 2012). On the contrary, lack of this proliferation of EGSCs during the embryo stage induces the opposite sex reversal in zebrafish and medaka (Kurokawa *et al*., 2007; Rodríguez-Marí *et al*., 2010), showing that this initial proliferation can be also involved in gonadal sex fate.

A second EGSC proliferation occurs prior to meiosis, following a special program of synchronous, successive and incomplete mitosis, generating interconnected daughter cells known as cysts or nests, which are surrounded by somatic cells (Saito *et al*., 2007). Following several rounds of mitosis, the cells enter meiosis to become oocytes in the ovary or spermatocytes in the testis (Lei and Allan C Spradling, 2013; Matova and Cooley, 2001; Tanaka, 2016). Cystic division is a highly conserved mechanism that precedes gametogenesis in both invertebrates and vertebrates (Amini *et al*., 2014; Hansen and Schedl, 2006; Hinnant *et al*., 2017; Lei and Allan C. Spradling, 2013; Quagio-Grassiotto *et al*., 2011). It has been well established in vertebrates that female germ cells enter meiosis earlier, while male germ cells are arrested and undergo meiosis at the onset of the prepubertal period (Elkouby and Mullins, 2017; Koubova *et al*., 2006; Lei and Allan C Spradling, 2013; Pepling, 2006). In medaka, this type of cystic cell division is known as proliferation type II, by which in females, the number of EGSCs increases dramatically during embryonic stages; whereas in males, this exponential proliferation to form cysts begins later, in the juvenile stage (Satoh and Egami, 1972; Tanaka, 2019). Despite the importance of cystic cell proliferation during early life to ensure reproductive success, this process is not fully understood at molecular level (Elkouby and Mullins, 2017).

In the search for genes that regulate EGSCs proliferation, *ndrg1* (n-myc downstream regulated gene 1) is one of the genes that was found to be upregulated in germ cell *foxl3* mutants, a switch gene involved in the germline sexual fate decision in medaka (Kikuchi *et al*., 2019). Interestingly, *Ndrg1* has been involved in the regulation of cell proliferation in different cancers (Chang *et al*., 2014; McCaig *et al*., 2011; Xi *et al*., 2017; Zhang *et al*., 2019). Moreover, this gene presents homologs from mammals (*Ndrg1*) to fish (*ndrg1*), *C*. *elegans* (ZK1073.1) to fruit fly (Dmel\MESK2), suggesting that NDRG1 is highly conserved among species (Kovacevic *et al*., 2013; Sun *et al*., 2013). *Ndrg1* has been demonstrated to inhibit the TGF-β signaling cascade, a very well-known pathway involved in cell proliferation and survival (Zhang *et al*., 2017), regulating the induction of epithelial-to-mesenchymal transition (EMT) in mammalian cells (Tojo et al., 2005; Chen *et al*., 2012; Jin *et al*., 2014), establishing *ndrg1* as a potent inhibitor of proliferation. However, its role in non-cancerous cells is not well established. In this study, we provide clear evidence of the important role of *ndrg1* in the regulation of EGSC cystic proliferation, with subsequent impact on reproductive success in medaka.

## MATERIALS AND METHODS

### Source of medaka

All experiments were performed with medaka (*Oryzias latipes*) (strain hi-medaka, ID: MT835) supplied by the National BioResource Project (NBRP) Medaka (www.shigen.nig.ac.jp/medaka/). Fish were maintained and fed following standard protocols for medaka (Kinoshita *et al*., 2012). Fish were handled in accordance with the Universities Federation for Animal Welfare Handbook on the Care and Management of Laboratory Animals (www.ufaw.org.uk) and internal institutional regulations. Fertilized medaka eggs were incubated in 60 mm Petri dishes with embryo medium (17 mM NaCl, 0.4 mM KCl, 0.27 mM CaCl2·2H2O and 0.66 mM MgSO4; pH 7) until hatching and subsequently in 2L water tanks to adults under a constant photoperiod (14L:10D) in a closed circulation water system at a controlled temperature (26 ± 0.5°C).

### Sample collection

Samplings were performed at different stages of medaka development: embryonic stages 35 and 39 (5- and 9-days post-fertilization, respectively) and juvenile-adult stages 10-, 20- and 80-days post-hatching (dph), according to a previous description of medaka development (Iwamatsu, 2004). These stages are important for medaka gonadal development. Stage 35 corresponds to a sexually undifferentiated gonad in both sexes, and only type I divisions (first mitotic proliferation). This stage precedes the beginning of type II division (cystic proliferation) in XX gonads (stage 36 - 39). Stage 39 corresponds to the maximum EGSC proliferation in XX embryos, and involves sexually dimorphic gonads, just at the time of hatching. At 10 dph, type II division is present only in XX gonads, while in XY gonads, it begins only at 20 dph. At 80 dph, animals have become sexually mature adults. For all sample collections, fish were euthanized by immersion in tricaine at 30±50 mg/L and processed according to the technique to be used. To determine genotypic sex, DNA from the tail and head of each animal was analyzed by PCR to determine presence of the *dmy/dmrt1bY* gene; ß-actin gene was used as a DNA loading control **(Table S1)** (Nanda *et al*., 2002). The PCR products were analyzed on 1% agarose gel.

### Whole-mount RNA *in situ* hybridization

Whole mount RNA *in situ* hybridizations were performed as previously described (Nakamura *et al*., 2006). Digoxigenin-labeled probes were synthesized from the full-length medaka cDNA of *ndrg1a* and *ndrg1b* using pGEM®-T Vector (Promega) linearized plasmid. Embryos at stage 35 were fixed overnight in 4% RNAse-free paraformaldehyde (PFA) at 4°C, permeabilized using 20 µg/µl proteinase K at room temperature (RT), and hybridized at 68°C overnight with either *ndrg1a* or *ndrg1b* digoxigenin (DIG)-labeled RNA probes. Hybridized probes were detected using an alkaline phosphatase–conjugated anti-digoxigenin antibody (1:2000; Roche) in the presence of nitro blue tetrazolium/5-bromo-4-chloro-3′-indolyphosphate substrates (Roche). Stained embryos and gonads were embedded in gelatin, cryostat sectioned at 14–16 µm thickness and photographed.

### RNA quantification by RT-qPCR

Embryos at stages 35 and 39 and juveniles at 10 and 20 dph were used for gene expression analysis. For this purpose, the gonadal region of the body was immediately frozen in liquid nitrogen and subsequently stored at −80°C until RNA extraction. Once the sex of each individual was known by PCR, total RNA was extracted from a pool of 5 embryos, or individually from juveniles, using 300 µL of TRIzol® Reagent (Life Technologies) following the manufacturer’s instructions. RNA from each sample (500 ng) was used to perform the cDNA synthesis, following previous studies (CastaΌeda Cortés *et al*., 2019). Real-time PCR primers are listed in Table S1. Gene-specific RT-PCR was performed using the SYBR green master mix (Applied Biosystem) and Agilent MX3005P Multiplex QPCR Real-time Thermal Cycler (Stratagene). The amplification protocol consisted of an initial cycle of 1 min at 95°C, followed by 10 s at 95°C and 30 s at 60°C for a total 45 cycles. The subsequent quantification method was performed using the 2-ΔΔCt method (threshold cycle; assets. thermofisher.com/TFS-Assets/LSG/manuals/cms_040980.pdf) and normalized against reference gene values for ribosomal protein L7 (*rpl7*) (Zhang and Hu, 2007) or elongation factor 1 alpha (*ef1α*) (Hirayama *et al*., 2006).

### TGF-β inhibitor treatment

To analyze whether TGF-β regulates *ndrg1b* expression, A83-01 (Sigma Aldrich, USA), a selective TGF-β inhibitor was used (Tojo et al. 2005). A83-01 inhibits the phosphorylation of Smad2, thereby blocking the signaling of TGF-β type I receptor (Tojo et al., 2005). A83-01 was first dissolved in DMSO (3 mM), and subsequently diluted in embryo medium (17 mM NaCl, 0.4 mM KCl, 0.27 mM CaCl2·2H2O, and 0.66 mM MgSO4; pH 7) to obtain a solution of 3 μM. Fertilized medaka eggs were incubated in 70 mm Petri dishes with or without 3µM A83-01 from stage 35 to stage 39 at 26°C **(Fig. S1)**. Freshly made dilutions were used and the medium was changed every 24h. As control, a group of embryos were incubated only with 0.1% DMSO. The concentration used in this study was based on Tojo *et al*. (2005). After the incubation period, embryos were collected at stage 39 for sex genotyping, RT-qPCR and cell quantification by histological analysis, as described in this article.

### Generation of *ndrg1b* mutants using CRISPR/Cas9-induced mutagenesis

CRISPR/Cas9 target sites were identified and designed using medaka genome reference available on the Ensembl genome database (ENSORLG00000004785), and the CCTop - CRISPR/Cas9 target online predictor (CCTop, http://crispr.cos.uni-heidelberg.de/) (Stemmer *et al*., 2015). Two sequences 5’GG-(N18)-NGG3’ in exon 6 of *ndrg1b* were identified. The selected target sites are shown in Fig. S1A and Fig. S2A. Each sgRNA was synthesized according a protocol previously established by Ansai and Kinoshita (2014). Briefly, the pair of inverse-complementary oligonucleotides were annealed and cloned into the pDR274 vector (Addgene #42250) in the BsaI cloning site. The modified pDR274 vectors were digested with DraI and each *ndrg1b* specific guide RNA (SgRNA_*Ndrg1b*) was transcribed using the MEGAshortscript T7 Transcription Kit (Thermo Fisher Scientific). The synthesized sgRNAs were purified by RNeasy Mini kit purification (Qiagen). For the Cas9-RNA in vitro synthesis, the pCS2-nCas9n vector (Addgene #47929) was linearized with NotI, and capped Cas9 RNA was transcribed with the mMessage mMachine SP6 Kit (Thermo Fisher) following the manufacturer’s instructions, and purified with the RNeasy Mini kit (Qiagen). Cas9-RNA (100 ng/µl) and SgRNA_*Ndrg1b* (50 ng µl) were co-injected into one-cell stage embryos as described previously (Kinoshita *et al*., 2000). Embryos injected only with *cas9* mRNA were used as controls (wild type: wt). Microinjections were performed with a Nanoject II Auto-Nanoliter Injector (Drummond Scientific) coupled to a stereomicroscope (Olympus).

### Biallelic CRISPR mutants screening and Off-target analysis

To analyze the efficiency and specificity of the CRISPR/Cas9 system, genomic DNA was extracted with alkaline lysis buffer using 3 dpf embryos, as described previously (Ansai and Kinoshita, 2014), DNA was used as a template for PCR to Heteroduplex mobility assay (HMA) using primers listed in Table S1. Electrophoresis was performed on 10% acrylamide gel and stained with ethidium bromide before the examination (Ota *et al*., 2013). Multiple heteroduplex bands shown by HMA in PCR amplicons from each injected embryo were quantified as embryos with biallelic mutations, whereas single bands were quantified as non-edited embryos. The mutation rate was calculated as the ratio of the number of multiple heteroduplex bands shown in PCR amplicons from each Cas9-sgRNA-injected embryo to the sum of all embryos injected multiplied by 100 (n=20/per sgRNA) (Ansai and Kinoshita, 2014). Sg1RNA_*ndrg1b* was selected for subsequent phenotypic analysis, due to its greater efficiency. Additionally, potential off-target sites in the medaka genome for Sg1RNA_*ndrg1b* were searched for by using the CCTop-CRISPR/Cas9 target online predictor (Stemmer *et al*., 2015). All potential off-target sites identified were analyzed by HMA using the primers listed in **Table S1**. Finally, the screening of indels was performed in F1 fish. Biallelic mutant adult (F0) medaka were mated with wild-type medaka. Genomic DNA was extracted from each F1 embryo for analysis of mutations by HMA, as described previously **(Table S1)**. Mutant alleles in each embryo were determined by sequencing of the *ndrg1b* genomic DNA region. All phenotypic analysis was performed using positive *ndrg1b* biallelic mutants F0 injected with Cas9 and Sg1RNA_*ndrg1b* (SgN1b) and Cas9-injected individuals as controls (wt).

### Immunofluorescence analysis

Gonadal regions from individuals at stage 39 were processed, fixed in Bouin’s solution, embedded in paraffin and sagittal sectioned at 5 µ m. Sections were washed with 0.1 M phosphate-buffered saline (PBS pH 7.4) and blocked in 0.1 M PBS containing 0.5% bovine serum albumin (Sigma-Aldrich) and 0.5% Triton X100 for 60 min before overnight incubation with primary antibody anti-OLVAS (1:200, rabbit, Abcam 209710) and anti-PCNA (1:200, mouse, Sigma P8825) at 4°C. After incubation, the sections were washed twice in PBS and incubated at RT for 90 min with Alexa Fluor 488-conjugated goat anti-rabbit IgG (ThermoFisher Scientific, A-11008) and 594-conjugated goat anti-mouse IgG (ThermoFisher Scientific, A32742) secondary antibodies at a dilution of 1:2000 in PBS. After incubation, sections were rinsed twice with PBS and mounted with Fluoromount mounting medium (Sigma-Aldrich) containing 4’,6-diamidino-2-phenylindole (DAPI, 5 µg/ml, Life Technologies).

### TUNEL assay

The presence of apoptosis at stage 39 was detected through the *In situ* Cell Death Detection Kit, Fluorescein (Roche). Samples fixed in Bouin’s solution, embedded in paraffin and sagittal sectioned at 5 µm were treated according to the manufacturer’s manual, with a step of permeabilization with 0.1% Triton X-100, 0.1% sodium citrate in PBS 1X solution. After labeling the reaction for 1 hour at 37°C, sections were immunostained with OLVAS, as described above.

### Histological analysis

Gonadal samples from juveniles at 20 (from whole body) and 80 dph (from gonadal tissue) were fixed in Bouin’s solution, embedded in paraffin, sectioned at 5 µ m, and subjected to standard Hematoxylin & Eosin staining. For the TGF-β inhibitor treatment, stage 39 individuals were fixed in 4% buffered glutaraldehyde, embedded in Technovit (7100-Heraeus Kulzer), sectioned at 5 μm thickness, and stained with 0.1% toluidine blue to quantify the germ cells using a light microscope (Leica DM6000 BD, Leica Microsystems).

### Cell quantification

All section photographs were taken using a ZEISS Axio Observer 7 Led colibri 7, Nikon Eclipse E600 and Nikon Digital Sight DSQi1Mc. Images were analyzed using FIJI software (https://imagej.nih.gov/ij/). Individual cells were counted manually with the Cell Counter plugin for FIJI. Type I and II germ cells were counted as previously described (Saito *et al*., 2007), where single isolated germ cells were counted as type I, while clusters with more than two germ cells were counted as germ cells undergoing type II division. Spermatogonia were counted based on the description by (Iwasaki *et al*., 2009)) and oocytes based on (Seki *et al*., 2004)) and Gay *et al*. (Gay *et al*., 2018).

### Sperm quantification

Sperm was collected from mutant and control adult males after immersion in tricaine at 30±50 mg/L. Each male was placed on a foam rubber holder with the belly facing up; the urogenital pore was dried and sperm was collected with a micropipette while gentle pressure was applied from bilateral abdominal toward the anal opening using fine forceps. Prior to the analysis, the sperm was diluted 5,000 times in cold Ringer’s solution (NaCl 0.75%, KCl 0.02%, CaCl_2_ 0.02%, NaHCO_3_ 0.002%) and kept at 4°C. Sperm counts were determined by hemocytometer (Bright-Line, Hausser Scientific) analysis under light microscope at 200X magnification, performing three counting replicates per male.

### Evaluation of reproductive success

To evaluate reproductive success, was used 3 categories of sexually mature couples: control (wild type female and male), mutant female (wild-type male and *ndrg1b* mutant female) and mutant male (wild type male and *ndrg1b* mutant male). Each couple were acclimatized for 5 days in a breeding tank to ensure and standardize their reproductive conditions. For 10 consecutive mornings, each couple’s fertilized and unfertilized eggs were carefully collected by hand, placed in a separate dish, and counted to calculate total spawned eggs, fertilization (number of fertilized eggs / number spawned eggs) and hatching (number of hatched larvae/number of fertilized eggs).

### Sexual behavior assay

Sexual behavior assay was performed over 10 days using 2 categories of sexually mature couples, control and mutant male, mentioned above, which were allowed to acclimatize for 5 days to ensure and standardize their reproductive conditions. On the remaining 5 days, the procedure was performed as previously described (Okuyama *et al*., 2014). Males and females were separated in the evening (6-7 PM) the day before the assay, using a transparent plastic cup with small holes for water exchange. The following morning, the mating pair were placed together in a single transparent tank, and their sexual behavior was recorded for 20 minutes using a digital video camera **(Fig. 6A)**. Water inflow to the tanks was stopped during recording. Females that did not spawn or spawned without any mating behavior were excluded from the analysis of parameters. The following behavioral parameters were calculated from the video recordings: latency until first courtship display, latency from the first courtship display to the wrapping that resulted in spawning; number of wrappings before and after spawning, the duration of wrapping, and within wrapping time, the duration of quivering and number of convulsions. Each action during sexual behavior was identified following the descriptions by Ono and Uematsu (1957) and Walter and Hamilton (Walter and Hamilton, 1970).

### Statistical analysis

Values are presented as mean ± standard error of the mean (SEM) for continuous variables and as percentages for categorical variables. Fold change and statistical analysis of RT-qPCR quantifications were performed using FgStatistics software (http://sites.google.com/site/fgStatistics/), based in the comparative gene expressions method (Pfaffl, 2001). Statistical analyses were performed by using GraphPad Prism (GraphPad Software, San Diego, CA). Continuous variables were compared between two groups by the unpaired two-tailed Student’s t-test. If the F-test indicated that the variance differed significantly between groups, Welch’s correction to the Student’s t-test was employed. For more than two groups, continuous variables were compared by one-way analysis of variance (ANOVA), followed by Dunnett’s post hoc test, for comparisons of experimental groups versus control group.). All differences were considered statistically significant when P<0.05.

## RESULTS

### Expression of *ndrg1* during gonadal development

To understand the role of *ndrg1* in cystic proliferation, the expression pattern of *ndrg1* during gonadal development was first determined. *Ndrg1* belongs to the n-myc downstream regulated gene 1 family, and in the medaka genome has two paralogs **(Fig. 1A)**, as a result of teleost-specific genome duplication (Kasahara *et al*., 2007): *ndrg1a* (ENSORLG00000003558) on chromosome 11 and *ndrg1b* (ENSORLG00000004785) on chromosome 16, with 60% identity. The *ndrg1a* transcripts were not detectable in the medaka gonads during embryo developmental stages by whole-mount RNA *in situ* hybridization **(Fig. 1B)**. In contrast, the medaka *ndrg1b* was specifically observed in gonads during embryo developmental stages **(Fig. 1C)**. Then, the transcript level of *ndrg1b* was quantified by qPCR at stage 35, when female (XX) and male (XY) gonads appear morphologically indistinguishable and germ cells only present type I proliferation. No difference for *ndrg1b* transcript levels was detected between female and male individuals **(Fig. 1D)**. Interestingly, differences between sexes were observed at stage 39, at 10 and 20 days post-hatching (dph) **(Fig**.**1E, F, G)**. At stage 39 and 10 dph, where female gonads have higher number of cysts compared to male gonads due to type II cystic division, the level of *ndgr1b* transcripts is higher in male than in female gonads **(Fig. 1E, F)**. The opposite sex transcriptional profile of *ndrg1b* was observed at 20 dph **(Fig. 1G)**, in which type II proliferation begins in male individuals. These results showed that transcription of *ndrg1b* is down-regulated during cystic proliferation of EGSCs in both sexes.

**Figure 1.**
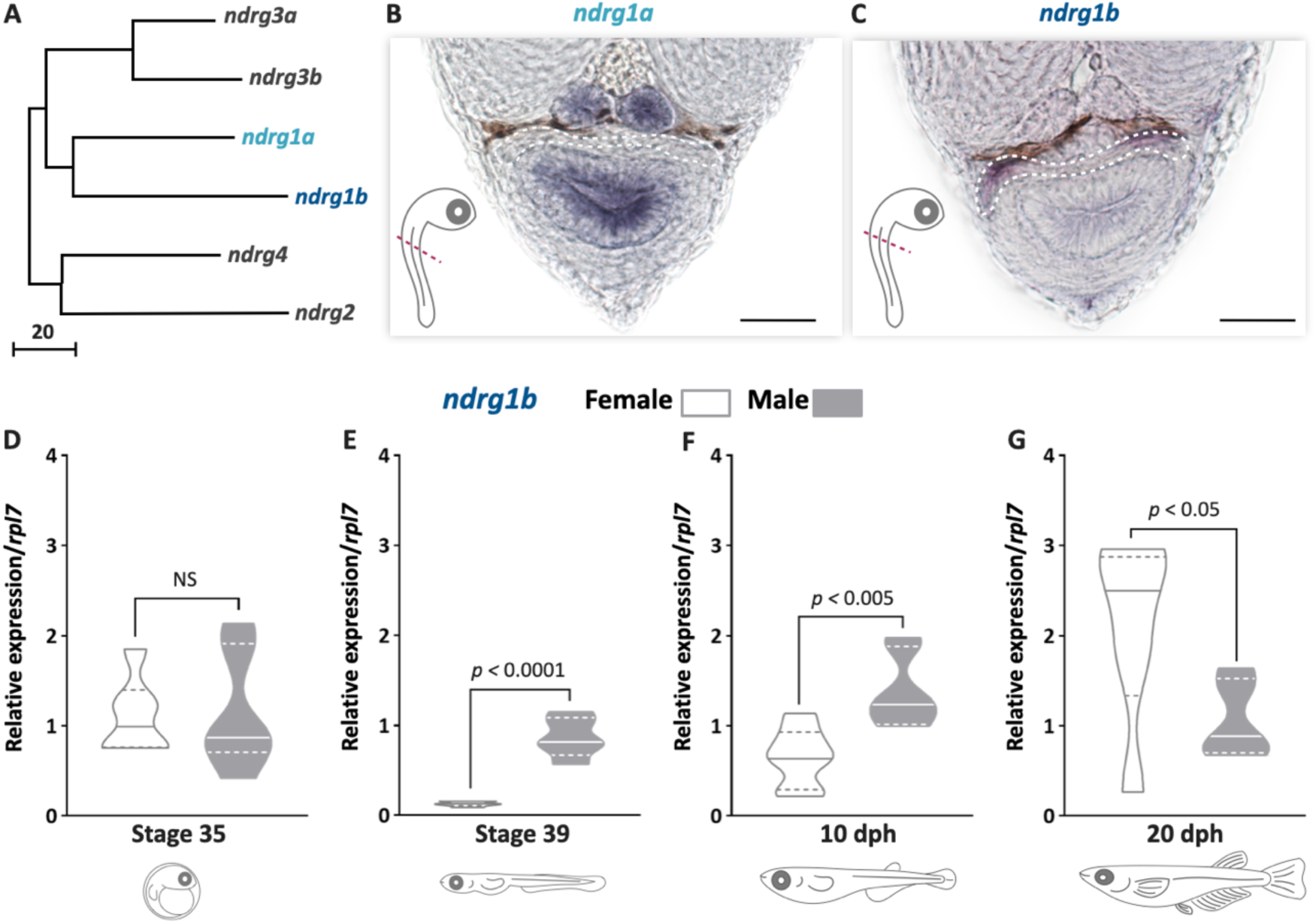
*ndrg1b* is down-regulated during cystic proliferation in gonadal development. Phylogenetic tree showing the relationships among the *ndrg* family components in medaka **(A)**, obtained using the neighbor-joining method and a bootstrap test (MEGA 7.0 software). The scale beneath the tree reflects sequence distances. ENSEMBL accession sequences are provided in Table S1. Transversal sections of gonadal region (red line) in whole embryos at stage 35 to detect *ndrg1a* **(B)** and *ndrg1b* **(C)** transcripts using *in situ* hybridization. Gonadal region is surrounded by white dotted line; scale bar represents 20µm. Transcript abundance levels of *ndrg1b* in different stages of gonadal development: 35 **(D)**, 39 **(E)**, 10 dph (days post hatching) **(F)** and 20 dph **(G)**. Quantification was performed using the 2^-ΔΔCt^ method and values were normalized to *rpl7*. Genotypic sex was determined by the presence/absence of the *dmy* gene; female (XX) and male (XY) are represented by empty bars or full bars, respectively. n= 5-6 pool per sex in stage 35 and 39 and n= 6-7 individuals per sex in 10 and 20 dph, P-values are indicated when transcript abundance between sexes at the same developmental stage differ statistically (P<0.05). NS, not statistically significant. Comparing relative gene expressions method (Pfaffl, 2001).

### TGF-β pathway during proliferation and its relation with *ndrg1b*

To analyze whether TGF-β signaling is involved in the regulation of cystic proliferation due the key role of the TGF-β superfamily in the regulation of type I proliferation in medaka (Imai *et al*., 2015; Nakamura *et al*., 2012), and the mediation of *ndrg1b* at it **(Fig. S1B)**, embryos of both sexes were exposed to A83-01 (TGF-β inhibitor), from stage 35 to stage 39 (**Fig. S1A**). When total of EGSCs was quantified, in exposed female no difference was observed, but an increased in EGSCs in exposed male was observed compared to control male (**Fig. S1C**). Additionally, no difference for *ndrg1b* mRNA levels was detected between control and treatment for female individuals (**Fig. S1D**). In contrast, male individuals showed increased levels of *ndrg1b* following TGF-β inhibitor treatment when compared to control (**Fig. S1D**), establishing that there is no negative feedback from *ndrg1b* via TGF-β. Overall, these results suggest that the TGF-β pathway is not involved in proliferation type II in medaka, and therefore it would not be regulated by *ndrg1b*. Further studies are therefore needed of other downstream *ndrg1b* pathways in the proliferation of germ cells during gonadal development, and elucidate the type of proliferation modified in A83-01 exposed male.

### Mutation of *ndrg1b* affects early gonadal development in both sexes

To explore the role of *ndrg1b* during gonadal development and cystic proliferation, firstly *ndrg1b* biallelic mutants (indels in F0 individuals genome created by injecting Cas9 and sgRNA) were generated using CRISPR/Cas9 technology. The mutagenesis efficiency for each sgRNA was analyzed by a heteroduplex mobility assay (HMA; **Fig. S2B, S3B**) (Ota *et al*., 2013), which reached 96.6% for sg1_*ndrg1b* and 80% for sg2_*ndrg1b* **(Fig. S2B, S3B)**, which is why sg1_*ndrg1b* was selected for subsequent analysis. In addition, biallelic mutants were mated with wild-type fish to generate F1 and then several fish were sequenced to confirm the presence of the indels **(Fig. S2C)**. Data indicate that most cells contained biallelic indels and, consequently, loss of function in *ndrg1b* mutants. None of the embryos analyzed presented indels at the off-target sites for the injected sgRNA of biallelic mutant **(Fig. S2D, E)**. In order to address the possible involvement of *ndrg1b* in sex development, biallelic mutants (F0) were analyzed.

Then, to analyze the phenotypic alterations on *ndrg1b* mutant gonad development, the number of EGSCs was counted using the germ cell marker *Olvas* at stage 39 **(Fig. 2A, B, C)**. The total number of EGSCs in male *sgN1b* was not affected with respect to the male wt individuals; however, the number was significantly reduced in female *sgN1b* compared to the male wt individuals **(Fig. 2D)**, resembling the number of male wt individuals. Moreover, when the number of type I and II germ cells in mutants were quantified, only type II cystic proliferation was altered in female mutants compared with wt male individuals **(Fig. 2F)**. Furthermore, the number of EGSCs inside each cyst in *sgN1b* was lower than in wt **(Fig. 2E)**.

**Figure 2.**
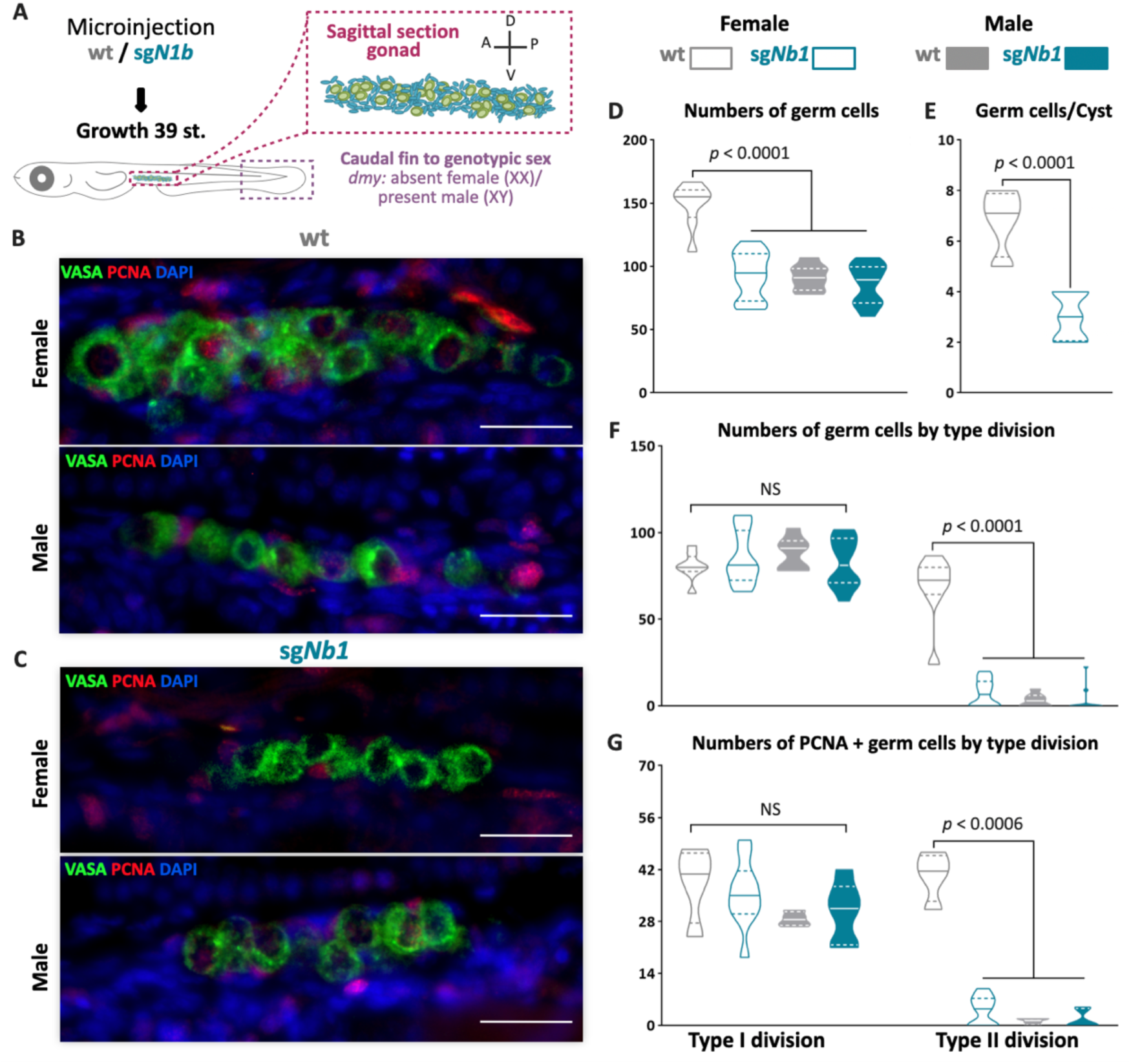
Mutation of *ndrg1b* affect gonadal development at stage 39. Schematic representation of the experimental procedure adopted to analyze the effect of *ndrg1b* loss on germ cell number and proliferation (A: anterior, P: posterior, D: dorsal and V: ventral) **(A)**. Fluorescent images of gonads in female or male embryos; wt =injected with *cas9* **(B)** and sg*Nb1*=injected cas9+sg1_*ndrg1b* **(C)**. EGSCs were stained with an anti-OLVAS antibody (green), nuclei were stained with DAPI (blue) and proliferating cells with PCNA antibody (red), scale bar represents 20µm. Total numbers of EGSCs **(D)**, number of EGSCs per cyst with type II division **(E)**, total numbers of germ cells with type I or type II division **(F)** and number of PCNA-positive type I and cystic type II germ cells relative to the total number of type I or type II germ cells, respectively **(G)**, in wt female or male embryos (represented by grey empty bars or full bars, respectively) and sg*N1b* female or male embryos (represented by cyan empty bars or full bars, respectively). Vertical bars indicate mean, with its respective s.e.m. n=8 per each wt group and n=12 per each sgN1b group. P-values are indicated when number of germ cells between groups differ statistically (P<0.05). NS, not statistically significant. Unpaired Student’s t-test. Dunnett’s post hoc test versus female wt.

Then we wondered whether EGSCs lose cystic proliferation or enter apoptosis in the *ndrg1b* mutants. The PCNA immunostaining **(Fig. 2B, C)** revealed that the number of proliferating type I and type II germ cells/total number of germ cell at stage 39 did not differ among female *sgN1b* and male gonads, but the number of proliferating type II germ cells was reduced in female *sgN1b* when compared to female wt **(Fig. 2G)**. In relation to the TUNEL assay, positive cells were not observed in any *sgN1b* or wt individuals **(Fig. S2A, B, C)**. Hence, female *ndrg1b* mutants presented less proliferating germ cells at stage 39. These results indicate that type II cystic proliferation is affected in *ndrg1b* mutants.

The effect of *ndrg1b* mutation was further analyzed during juvenile development in both sexes at 20 dph, which is the time when type II cystic division begins in males. Histological examination showed ovaries with apparently normal development **(Fig 3B)**; however, there is a decrease in the number of oogonia and pre-vitellogenic oocytes compared to wt individuals **(Fig 3D, E)**. With regard to the males, the mutant showed a significant reduction of cysts and germ cells within the cyst in relation to wt males **(Fig 3F, G)**. This indicates that germ cell number is affected in *ndrg1b* mutants for both sexes at times that are key to the commitment of type II cyst division.

**Figure 3.**
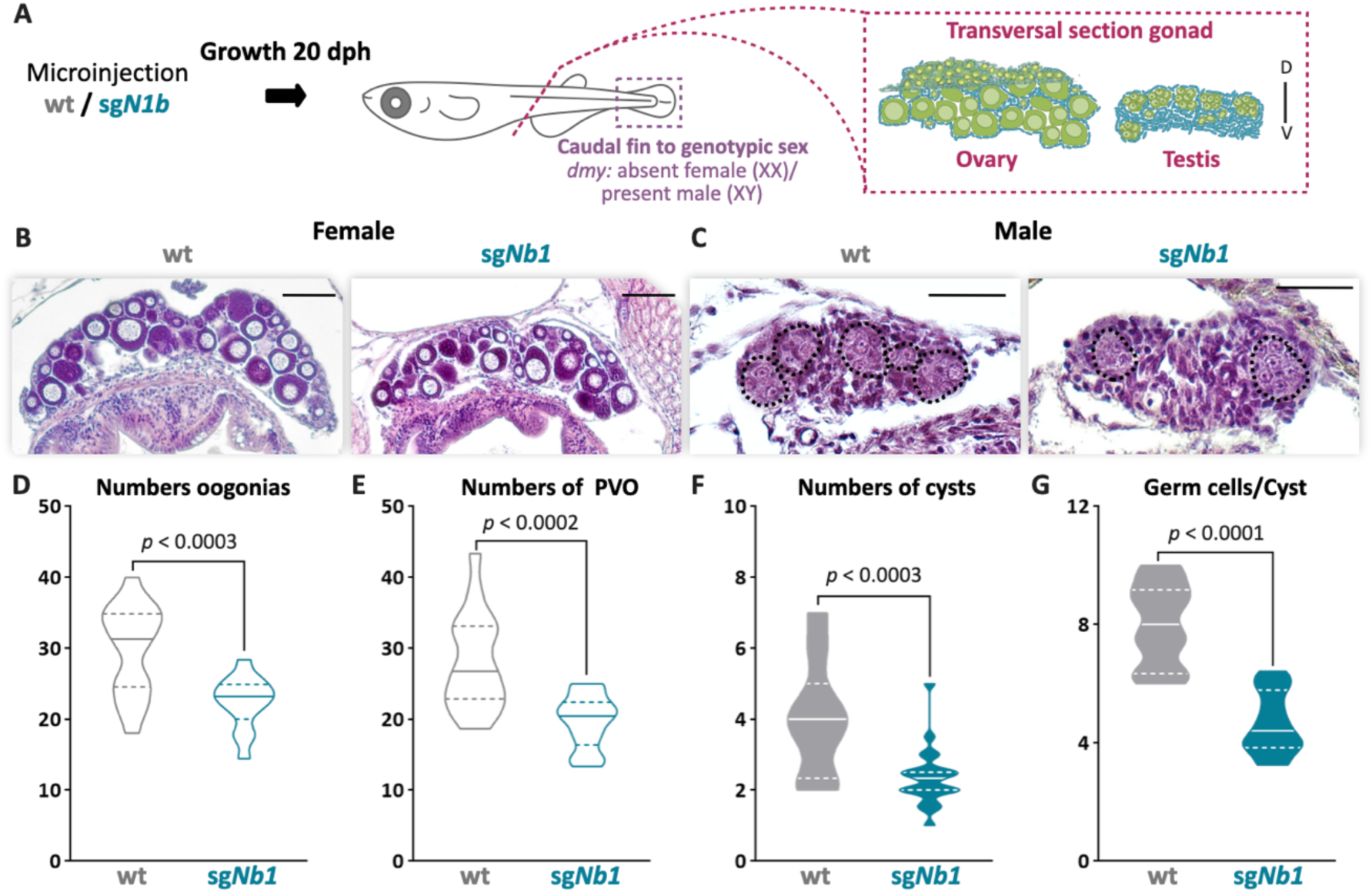
The mutation of *ndrg1b* affects gonadal development at 20 dph. Schematic representation of the experimental procedure adopted to analyze the effect of *ndrg1b* mutation on gonadal histology at 20 days post-hatching (dph, D: dorsal, V: ventral) **(A)**. Histological transversal sections of gonads wt= injected with *cas9* or sg*Nb1*=injected with cas9+sg1_*ndrg1b* from individuals for XX **(B)** and XY **(C)**. In the testis, each cyst of germ cells (spermatogonia-like cells) is encircled by a black dotted line. Scale bars in B and C represent 50µm. Total number of oogonia **(D)**, number of pre-vitellogenic oocytes (PVO) **(E)**, number of spermatogonial cysts **(F)**, and number of spermatogonia per cyst **(G)**, in Wt female or male embryos (represented by grey empty bars or full bars, respectively) and sg*N1b* female or male embryos (represented by cyan empty bars or full bars, respectively). n=16 per female groups and n=20 per male groups. P-values are indicated when number of germ cells between groups differs statistically (P<0.05). NS, not statistically significant. Unpaired Student’s t-test.

In addition, we performed an extra check of the efficacy of the loss of function of *ndrg1b* with the *sgN1b* guide, by analyzing the phenotype of loss of cystic proliferation in individuals injected with a second RNA guide, *sg2_ndrg1b*. We observed the same results as the *sgN1b* biallelic mutants (**Fig. S3C, D**). These results reinforce the results obtained with the biallelic mutants of *ndrg1b*.

### Mutation of *ndrg1b* alters gametogenesis and reproductive success of both sexes in adulthood

To evaluate whether impaired type II cystic proliferation can affect gametogenesis and reproductive success, paired adult male and female wt and *ndrg1b* mutants were analyzed for 10 days, and then their body and gonadal morphology was analyzed according to different reproductive parameters (**Fig. 4A**). Regarding the daily frequency of spawning, the *ndrg1b* mutant females spawned fewer eggs than wt females (**Fig. 4B**), although fertilization rate was normal (**Fig. 4C**). Surprisingly, the wt females paired with male *sgN1b* also spawned fewer oocytes (**Fig. 4B**), and in *sgN1b* males paired with wt females, the fertilization rate declined (**Fig. 4C**). Moreover, hatching rate was lower for the progeny of male *sgN1b* and wt female than for the progeny of female *sgN1b* and wt male (**Fig. 4D**).

**Figure 4.**
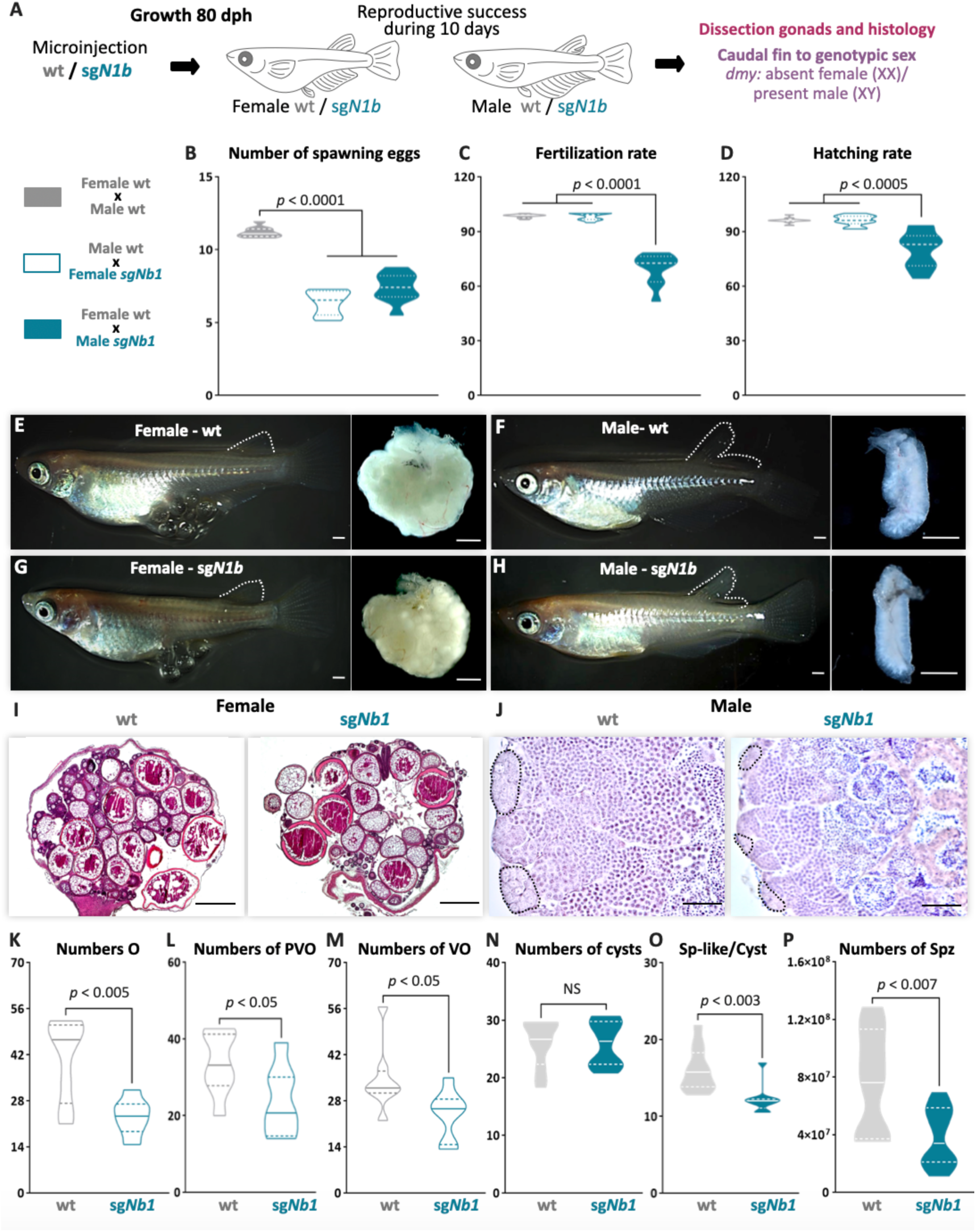
Mutation of *ndrg1b* alters the gametogenesis and reproduction success of both sexes in adulthood. Schematic representation of the experimental procedure adopted to analyze the effect of *ndrg1b* loss of function in the reproductive success at 80 days post-hatching (dph) **(A)**. Reproductive success was evaluated by crossing the wt males with *sgN1b* females (cyan full), wt females with *sgN1b* males (cyan empty), and as a control, wt females with wt males (grey). For all crossings, we quantified the total number of eggs spawned **(B)**, percentage of fertilization **(C)**, and percentage of hatching **(D)** of the total eggs spawned. The biallelic mutants of *ndrg1b* did not exhibit morphological sexual differences: adult females with their ovaries, wt **(E)** and sg*Nb1* **(G)**, and adult males with their testis, wt **(F)** and sg*Nb1* **(H)**. The secondary sexual characteristics are indicated by the separation of hindmost rays from other rays in the dorsal fin. Scale bars are 1mm. Histological transversal sections of wt and sg*Nb1* gonads from XX **(I)** or male **(J)** individuals. In the testis, each cyst of germ cells (spermatogonia-like cells) is encircled by a black dotted line. Scale bars represent 500 µm (I) and 50 µm (J). Number of oogonia (O) **(K)**, Number of pre-vitellogenic oocytes (PVO) **(L)**, number of vitellogenic oocytes (VO) **(M)** number of spermatogonial cysts **(N)**, Number of spermatogonia per cyst **(O)**, and number of spermatozoa **(P)** from wt female or male individuals (represented by grey empty bars or full bars, respectively) and sg*N1b* female or male individuals (represented by cyan empty bars or full bars, respectively). n=10 per group. P-values are indicated when groups differ statistically (P<0.05). NS, not statistically significant. Dunnett’s post hoc test versus control couples of wt female and male. Unpaired Student’s t-test.

Medaka adults display easily distinguishable secondary sex characteristics, e.g. the hindmost rays are separated from other rays in the dorsal fin in the male, but linked together in female. All *ndrg1b* mutants invariably displayed typical female and male secondary sex characteristics **(Fig. 4E, F, G, H)**. A balanced sex ratio was observed in both *ndrg1b* mutants and wt individuals **(Table S2)**. Histological examination of females showed ovaries with apparently normal development **(Fig. 4I)**, but there were significantly fewer oogonia, pre-vitellogenic and vitellogenic oocytes at the different stages in *ndrg1b* mutants than in wt individuals **(Fig. 4K, L, M)**. With respect to the histological examination of testis, all stages of spermatogenesis were observed in *sgN1b* **(Fig. 4J)**. Spermatogonia are surrounded by Sertoli cells at the distal end of the lobule, which are responsible for constantly supplying new germ cells by proliferation (Iwai *et al*., 2006). When these cells were counted manually, the number of cysts containing spermatogonia remained unchanged **(Fig. 4N)**; however, the number of spermatogonia within each cyst was lower in *sgN1b* than in wt individuals **(Fig 4O)** and the total number of spermatozoa was lower in *sgN1b* testes than in wt testes **(Fig. 4P)**. Collectively, these results indicate that alterations of cystic proliferation, with the concomitant lower sperm count and fewer oocytes, have direct influence on reproduction. Interestingly, these alterations do not change the general gonadal morphology or secondary sex characteristics in either sex, or the sex ratio.

### Mutation of *ndrg1b* disturbs the mating behavior of males

To explore a possible cause of lower spawning of control females paired with male *ndrg1b* mutants, our next step explored the sexual behavior of these couples. Fortunately, the sexual behavior of medaka consists of a sequence of actions that are easily quantified (Hiraki-Kajiyama *et al*., 2019; Ono and Uematsu, 1957; Walter and Hamilton, 1970). Briefly, the sequences begin with the male approaching and following the female closely. The male then performs a courtship display, in which the male swims quickly in a circular pattern in front of the female. If the female is receptive, the male grasps her with his dorsal and anal fins (termed ‘wrapping’), they quiver together (termed ‘quivering’), and the male make several gentle taps, like “convulsions” (termed “convulsions”), and then oocytes and sperm are released. In general, all couples displayed the usual actions of sexual behavior **(Fig. 5B-I)**, with differences observed only in the actions related to mating behavior, particularly in the wrapping actions related to the stimulation of the female spawning, the duration of quivering and number of convulsions, which were significantly fewer in *sgN1b* males paired with wt females than in control couples **(Fig. 5H, I)**. Collectively, these results indicate that alterations of cystic proliferation, with the concomitant lower sperm count, have direct influence on reproduction and mating behavior.

**Figure 5.**
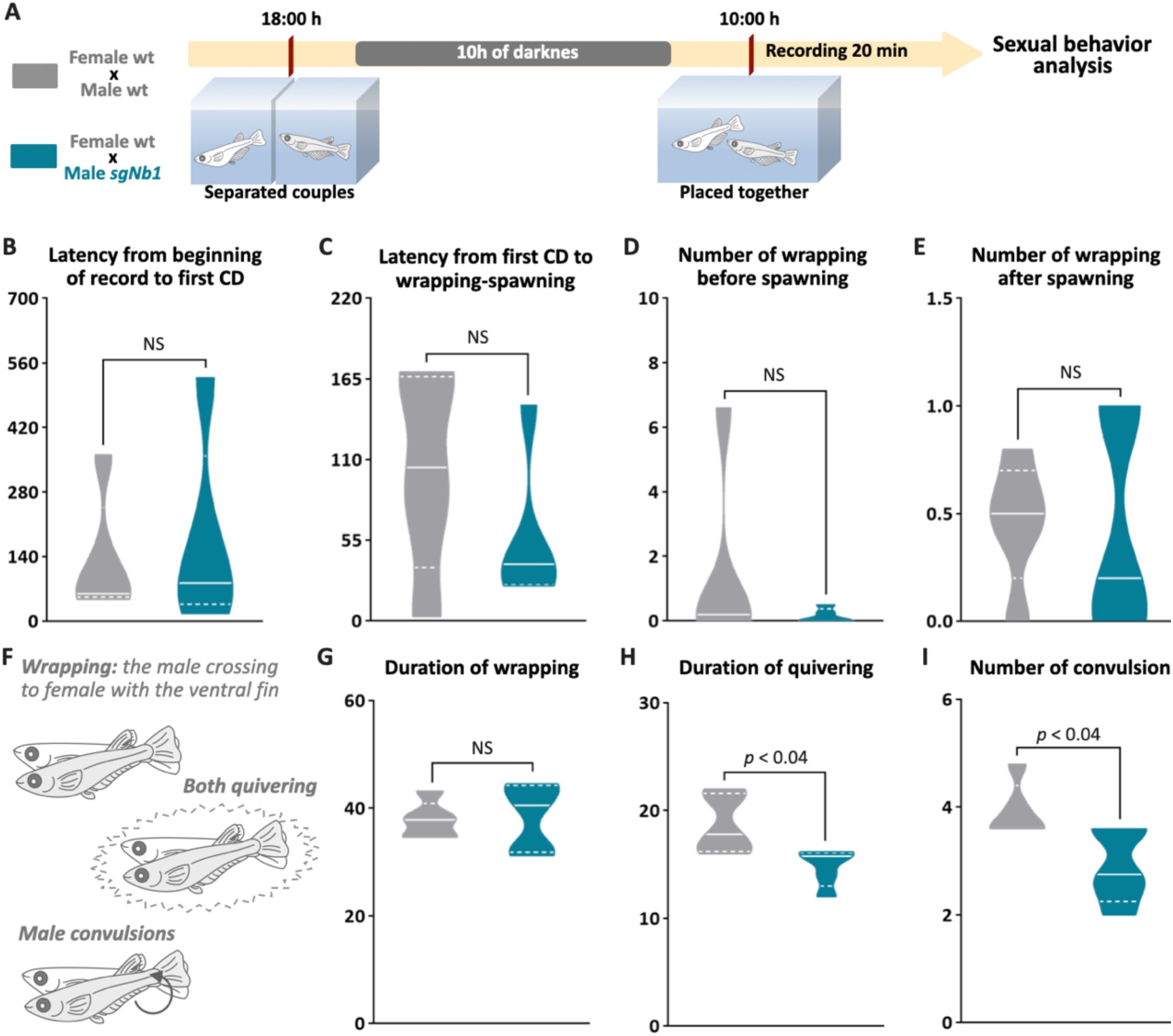
Reduction of cystic proliferation in *ndrg1b* mutants decreases male sexual behavior, with a decrease of mating vigor. Schematic representation of the experimental procedure adopted to analyze reproductive behavior for 5 days; five couples per group were analyzed **(A)**. Couples of wt individuals (gray) or *sgN1b* males with wt females (cyan), were separated in the evening (18:00–19:00) the day before the assay, using a transparent plastic tank with small holes for water exchange. The following morning, each mating pair was placed together in a single tank and sexual behavior was recorded for 20 minutes. Each video was analyzed to determinate the latency from beginning of recording to the first courtship display **(B)**, latency from first courtship display to the wrapping that resulted in spawning **(C)**, number of wrappings of 2 seconds of duration before **(D)** and after **(E)** spawning, and specific wrapping actions **(F)**, such as duration of wrapping **(G)**, duration of quivering **(H)** and number of convulsions **(I)**. Representative videos of wrapping actions in couples of wt female and male (https://www.youtube.com/watch?v=KrI8t90_tMA&feature=youtu.be) and male sgN1b with female wt (https://www.youtube.com/watch?v=MCjhYG7lfwM&feature=youtu.be) n=5 per group. P-values are indicated when number of germ cells between groups differ statistically (P<0.05). NS, not statistically significant. Unpaired Student’s t-test.

## DISCUSSION

*Ndrg1* homologs have been found in different species of invertebrates and vertebrates (Fang *et al*., 2014; Kovacevic *et al*., 2013; Sun *et al*., 2013). In mammals, it has been well established that *Ndrg1* exerts an inhibitory effect on cancer cell proliferation (Chang *et al*., 2014; McCaig *et al*., 2011; Xi *et al*., 2017; Zhang *et al*., 2019). Additionally, several studies have shown that *Ndrg1* is expressed in the gonads of humans, mice, sheep and medaka (Burl *et al*., 2018; Kikuchi *et al*., 2019; Lachat *et al*., 2002; Qu *et al*., 2019; Stévant and Nef, 2019), however its role in gonadal function and reproduction is still unknown. In medaka, there are two paralogs of *ndrg1*: *ndrg1a* and *ndrg1b*, although only *ndrg1b* is expressed in the gonad. Interestingly, *ndrg1b* expression is opposite to the EGSC cystic proliferation in both sexes. Present from invertebrates to vertebrates, cystic division is a highly conserved mechanism that precedes gametogenesis and is crucial to reproductive success. However, its molecular regulation remains unknown (DiNardo *et al*., 2011; Hinnant *et al*., 2017; Narbonne-Reveau *et al*., 2006). In the present study, we generated *ndrg1b* F0 mutants by using CRISPR/Cas9 technology, and showed that the lack of *ndrg1b* leads to a reduction in the total number of germ cells by inhibiting cystic proliferation in both sexes during early gonadal development and adulthood. This is evidence that *ndrg1b* is likely involved in the molecular regulation of cystic division in medaka. Gametogenesis commences within the spermatocysts or cysts in males and within germline cradles in females, from germline stem cell precursors called spermatogonia or oogonia, respectively (Nishimura *et al*., 2016). Histological examination of adult gonads showed that medaka *ndrg1b* mutants displayed typical female and male gonadal structure with all the different stages of germ cells; however, they had a lower production of gametes. In this regard, although a previous study has shown that type I proliferation is important to germ stem cell number in adult medaka gonads (Morinaga *et al*., 2007), here we showed that type II proliferation (cystic) is also crucial to gamete production, and consequently, to related reproductive parameters such as the number of spawned eggs, spermatozoa and fertilization rate.

Additionally, we observed that the reduction of cystic proliferation in *ndrg1b* mutants did not induce any sex reversal or modification in the sex ratio. These observations are in agreement with Nishimura et al. (2018) in *figla, meioC* and *dazl* mutants in medaka, in which follicle formation is disrupted, germ cells are unable to commit to gametogenesis and germ cells do not develop into EGSCs, respectively. All these mutants exhibited female secondary characteristics and ovary structures, demonstrating that the mechanism underlying sex-reversal is manifested before germ cells exit from the status of stem cells/gonocytes and enter the gametogenic program (Nishimura *et al*., 2018). In contrast, medaka mutants which had an effect on proliferation type I, like the germ cell-deficient *cxcr4* (Kurokawa *et al*., 2007) and also loss of *amhrII* (TGF-β pathway member) that display hyperproliferative germ cells (Morinaga *et al*., 2007; Nakamura *et al*., 2012), showed sex reversal. Interestingly, our results showed that the TGF-β pathway is not involved in the cystic division, and the increase proliferation in males is due to inhibition of TGF-β members involved in controlling germ cell proliferation type I, such as *amh*. Further studies are therefore necessary to elucidate the regulatory pathways of this proliferation type.

An interesting observation in the present work was the lower spawn by normal females paired with *ndrg1b* mutant males. Have been established that sexual behavior is linked to reproductive success (Oshima *et al*., 2003) and fortunately, the sexual behavior of medaka consists of a sequence of several actions which are easily quantified (Ono and Uematsu, 1957; Walter and Hamilton, 1970). We specifically observed differences in male mating behavior related with stimulation of female spawning and fertilization, such as the duration of quivering and number of convulsions; actions that seems to be conserved in vertebrates, because the ejaculation in male rat is accompanied by severe vibration and significant abdominal contraction (Qin *et al*., 2017). Sexual behavior has been found to be strongly controlled by the brain (Hiraki-Kajiyama *et al*., 2019; Mitchell *et al*., 2020; Yang and Shah, 2014), and levels of sex steroid hormones (Hiraki-Kajiyama *et al*., 2019; Munakata and Kobayashi, 2010). Here, it is important to note that in medaka, the secondary sex characters are induced by sex steroid hormones (Kikuchi *et al*., 2019; Sato *et al*., 2008). Since *ndrg1b* mutants presented both male and female secondary sex characters, it is suggested that sex steroid synthesis is not impaired in these animals. Regarding sexual behavior, in studies where the regulation of sex hormones was altered, changes in sexual behavior were primarily related to courtship, such as the frequencies of following, dancing, and latency from the first courtship (Hiraki-Kajiyama *et al*., 2019; Oshima *et al*., 2003). Additionally, when the brain regulators were altered, such as arginine-vasotocin (Yokoi *et al*., 2015), oxytocin ligand (oxt) (Yokoi *et al*., 2020) and TN-GnRH3 (Okuyama *et al*., 2014), changes in courtship display and sexual motivation, latency to mating and number of courtship displays, and latency to mating, were observed, respectively. None of these courtship actions changed in the male *ndrg1b* mutants, leading us to consider whether the specific mating actions of quivering and convulsions are exclusively related to the lower number of spermatogonia and spermatozoa in males. To our knowledge, only one other study has observed this loss of mating vigor in medaka. In it, successive multiple mating-induced depletion of sperm reserves decreased fertilization and mating rate, and even females responded by reducing clutch size (Weir and Grant, 2010). In this respect, this association warrants further investigation in vertebrates including brain, endocrine and gonadal features.

## CONCLUSION

Taken together, our data show that mutations of *ndrg1b* lead to disruption of EGSC cystic proliferation with a subsequent reduction in the number of gametes and an unexpected change in mating behavior of medaka. Altogether, these data are significant for three main reasons: 1) we showed that *ndrg1b* is involved in the regulation of cystic proliferation during different stages of medaka development (embryo, juvenile and adult) in a TGF-β-independent manner; 2) we demonstrated that mutation of *ndrg1b* leads to changes in number of gametes, decreasing reproductive success for both sexes; and 3) *ndrg1b* mutation affected male mating behaviors in medaka.

## CREDIT AUTHOR STATEMENT

LFAP: Conceptualization, Methodology, Writing-Original draft preparation, Visualization; DCCC: Methodology and Visualization; IFR: Methodology; RHN: Methodology and Writing-Reviewing and Editing, JIF: Funding acquisition, Conceptualization, Investigation, Supervision; Writing-Reviewing and Editing.

## ACKNOWLEDGEMENTS

We thank Technician Gabriela C. López and Dr. Leandro A. Miranda (INTECH) for helping with histological preparations. We also thank Dr. Tania Rodriguez for technical support and Technician Javier Herdman (INTECH) for fish handling. We are grateful to NBRP Medaka (https://shigen.nig.ac.jp/medaka/) for providing HNI (Strain ID: MT835).

