## Supplementary Figures and Tables for "Cystic proliferation of embryonic germ stem cells is necessary to reproductive success and normal mating behavior in medaka"

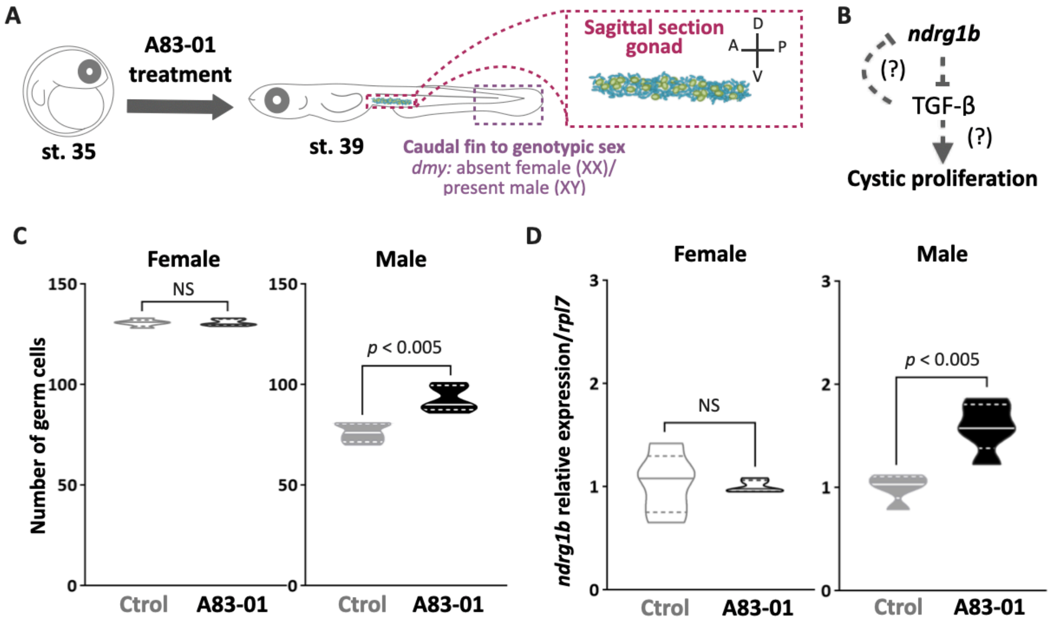


**Figure S1. TGF-β is not involved in the *ndrg1b*-mediated cystic proliferation in females during early gonadal development.** Schematic representation of the experimental procedure adopted for TGF-β inhibitor (A83-01) treatment **(A)**. Proposed hypothesis **(B)**. Total number of EGSCs in XX group and XY group **(C)** in embryos at stage 39 treated with A83-01**.** n= 5 pools per group, P-values are indicated when transcript abundance and number of germ cells between treated group and untreated group (Ctrol) of the same sex differ statistically ($P<0.05$). Transcript abundance levels of *ndrg1b* in XX and XY **(D)** embryos at stage 39 treated with A83-01. Quantification was performed using the $2^{-\Delta\Delta Ct}$ method and values were normalized to *rpl7* (n= 5 pools per group). Genotypic sex was determined by the presence/absence of the *dmy* gene; female (XX) and male (XY) are represented by empty bars or full bars, respectively. NS, not statistically significant. Comparing relative gene expressions method (Pfaffl, 2001) to transcript abundance and unpaired Student's t-test to germ cell number.

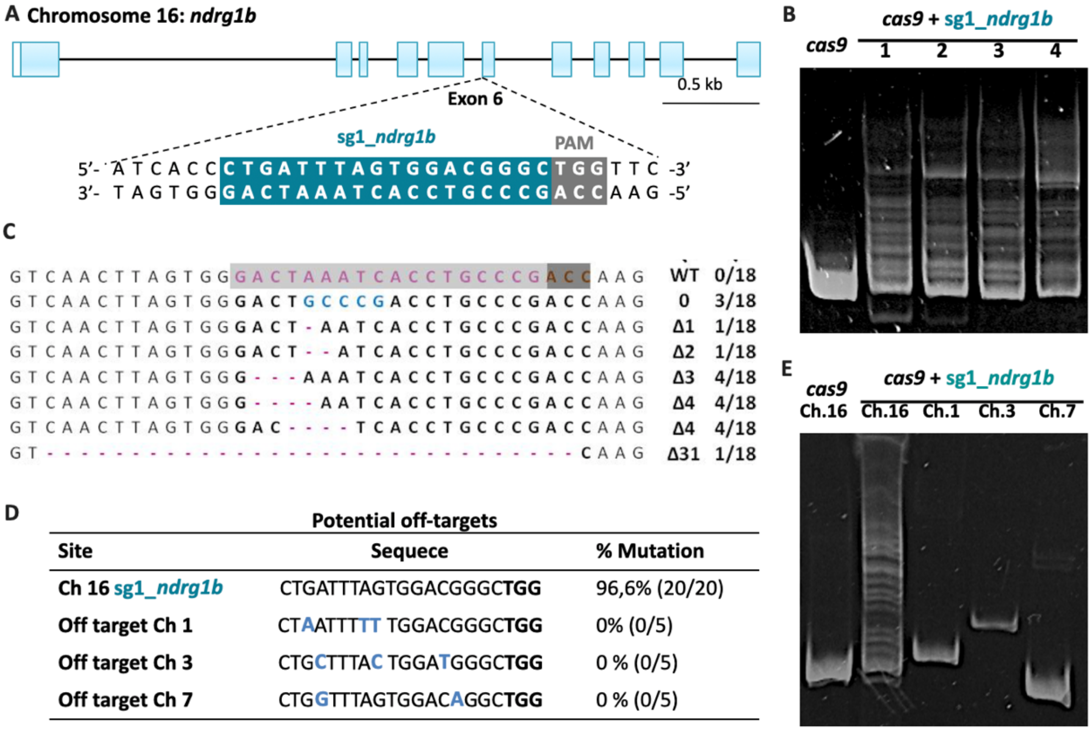


**Figure S2. CRISPR/Cas9: sg1_*ndrg1b* design, heteroduplex mobility assay (HMA), efficiency and potential off-targets*.*** Schematic representation of the genomic structure of *ndrg1b*, coding exon regions are shown as blue solid boxes, the targeting sequence of sg1_*ndrg1b* is indicated by cyan box, adjacent to NGG protospacer adjacent motif (PAM) sequence in gray box **(A)**. Images of heteroduplex mobility assay (HMA) to cas9 injected embryo (wild type, single band) and *cas9* + sg1_*ndrg1b* injected embryos (1-4, multiple bands) **(B)**. Subcloned sequences observed in the *cas9* and *cas9*+sg1_*ndrg1b embryos at F1* **(C)**, the targeting sequence of the sg1_*ndrg1b* is indicated by a cyan box, adjacent to PAM sequence in dark gray box. Blue letters indicate the identified insertion and purple dashes indicate the different deletions. The size of *indels* is shown to the right of each mutated sequence. Numbers on the right edge indicate the numbers of mutated clones identified from all analyzed clones from each embryo. Table indicates the target and potential off-target loci identified with respective chromosome (Ch), sequence, percentage of biallelic mutation detected **(D)**, the number of embryos with biallelic mutations/total eggs injected are shown in parentheses. Sequences of the off-target primers used for HMA are listed in Table S1. Image of heteroduplex mobility assay (HMA) for detecting off-target alterations **(E)**, potential off-target loci for sg1_*ndrg1b* were analyzed using genomic DNA extracted from five embryos co-injected with 100 ng/mL of *cas9* and 25 ng/mL of sg1_*ndrg1b*.

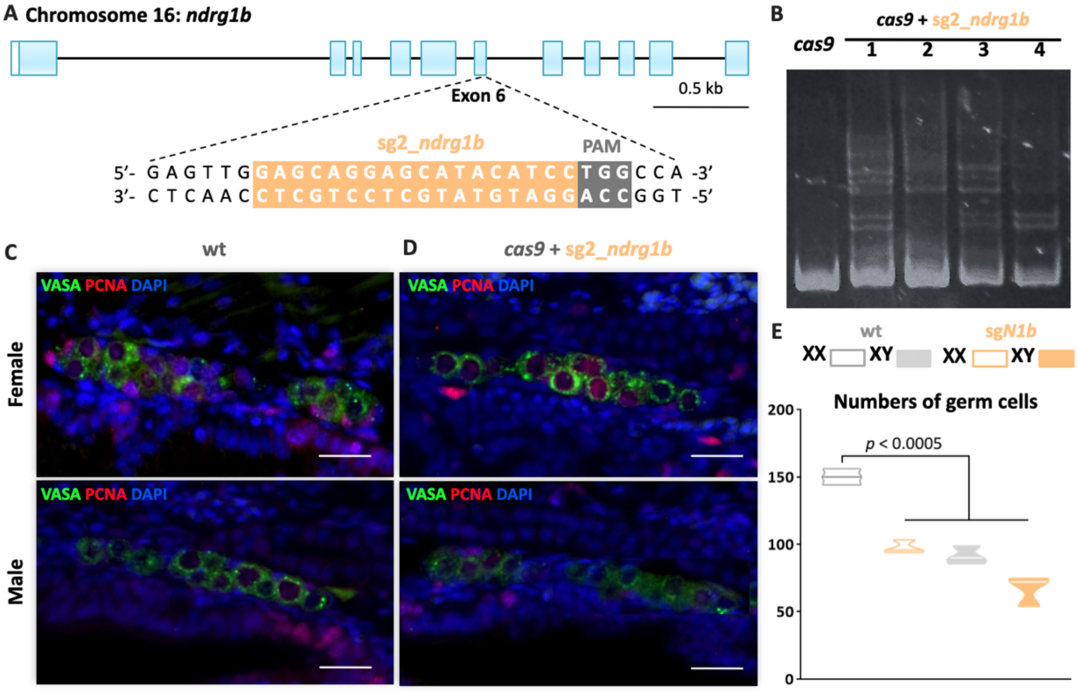


**Figure S3. Corroboration with a second RNA guide (sg2_*ndrg1b*) the specificity of CRISPR/Cas9 methodology to mutate *ndrg1b*.** Schematic representation of the genomic structure of *ndrg1b*, coding exon regions are shown as blue solid boxes. The targeting sequence of sg2_*ndrg1b* is indicated by orange box, adjacent to NGG protospacer adjacent motif (PAM) sequence in gray box **(A)**. Images of heteroduplex mobility assay (HMA) to cas9-injected embryo (wild type, single band) and *cas9*+sg2_*ndrg1b*-injected embryos (1-4, multiple bands) **(B).** Fluorescent sagittal images of gonads from embryos XX or XY injected with *cas9* **(C)** or cas9 + sg2_*ndrg1b* **(C)** at stage 39. Germ cells were immunostained using an anti-OLVAS antibody (green), nuclei were stained with DAPI (blue) and proliferating cells with anti-PCNA antibody (red). Scale bars represent 50µm. The total numbers of germ cells in cas9 and sg2_*ndrg1b* female or male embryos **(E)**. n= 3 per group. P-value is indicated when number of germ cells between groups differs statistically (P<0.05). Dunnett's post hoc test versus female wt.

**A**

| | % TUNEL + EGSCs | |
|---|---|---|
| | *cas9* | sg*N1b* |
| **Female (XX)** | 0 | 0 |
| **Male (XY)** | 0 | 0 |

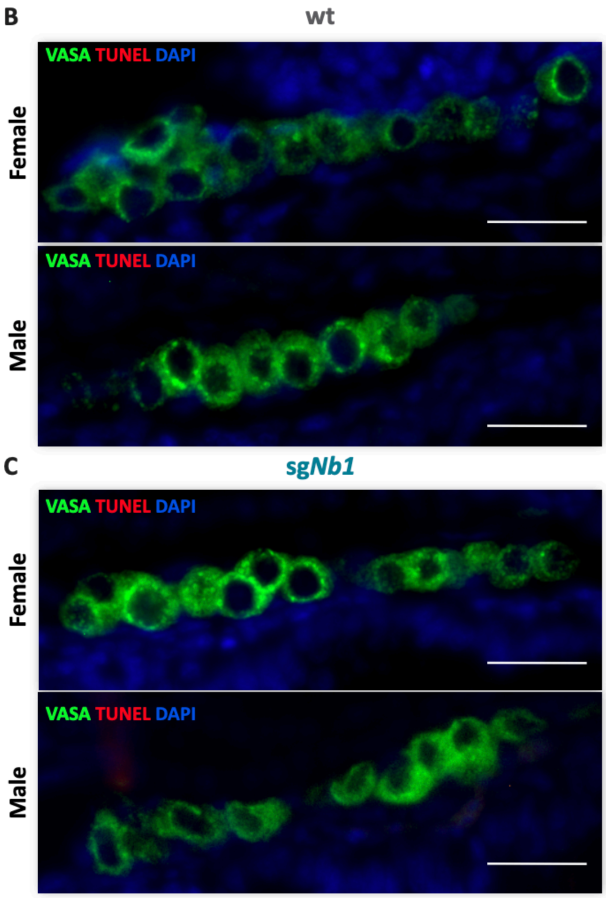


**Figure S4. Apoptosis of EGSCs at stage 39.** Percentage of TUNEL-positive EGSCs at stage 39 were determined using 10 sagittal sections per individual (n=4) **(A)**. Fluorescent images of gonads from female or male embryos injected with *cas9* **(B)** or sg*Nb1* (cas9+sg1_*ndrg1b*) **(C)** at stage 39. Germ cells were immunostained using an anti-

OLVAS antibody (green), nuclei were stained with DAPI (blue) and TUNEL positive cells (red). Scale bars represent 20μm.

| Gene symbol | Accession Number | primer sequence (5´- 3´) | Experiment |
|---|---|---|---|
| ***ndrg1a*** | ENSORLG00000003558 | **Fw:** GACGATATCCAGGTTGTCGAGTCC **Rv:** CGATGATATGGTTGCGATATCTCGC | **WMISH** |
| ***ndrg1b*** | ENSORLG00000004785 | **Fw:** CATGTTGAGGCTCCAGGACAAC **Rv:** CTGCAGCTCGTTGGTATGTGAG | |
| ***ndrg1b*** | ENSORLG00000004785 | **Fw:** ATGTCAACCCCAATGCTGAG **Rv:** CGTTGGACTGGTTCATGGTT | **RT-qPCR** |
| ***RPL7*** | ENSORLG00000007967.2 | **Fw:** CGCCAGATCTTCAACGGTGTAT **Rv:** AGGCTCAGCAATCCTCAGCAT | |
| ***ef1*** **α** | ENSORLG00000007614 | **Fw:** GGAGGCCAGCGACAAGATGAGC. **Rv:** ACACGGCCGACAGGGACAGTTC | |
| ***dmy*** | ENSORLG00000020486 | **Fw:** CAACTTTGTCCAAACTCTGA **Rv:** TGATGCAGCATTTTGACACATTTA | **Sexing PCR** |
| ***B-actin*** | ENSORLG0000001367 | **Fw:** GGATGACATGGAGAAGATCTGG **Rv:** ATGGTGATGACCTGTCCGTC-3' | |
| ***ndrg1b*** | ENSORLG00000004785 | **Fw:** CTTGACATGCCTTTATCTAGAGAC **Rv:** CATTTCTGTCACCAGCTTAAG | **HMA** |
| ***Chr1*** | chr1:5601523-5601805 | **Fw:** GGGTGAATGTTGCAGAAGTTG **Rv:** GGACTGTTGGAATGTGGGTG | **Off-target HMA** |
| ***Chr3*** | chr3:4654886+4655208 | **Fw:** CTCTGTGGTCCAGTACCAACTG. **Rv:** GGTGAGTTGATAATGCGGTC | |
| ***Chr7*** | chr7:21163167+21163418 | **Fw:** GGAGCCGCCTGTTAGCTTC **Rv:** CTGGCTCGTGTGACACATACG | |

**Table S1.** Primers sequences, ENSEMBL accession numbers and respective references were added to each gene.

| Sex ratio | | | | | |
|---|---|---|---|---|---|
| | wt | | sg*N1b* | | % sex reversal |
| | Testis | Ovary | Testis | Ovary | |
| **Female (XX)** | 0 | 26 | 2 | 21 | 8,70 |
| **Male (XY)** | 30 | 0 | 28 | 1 | 3,45 |

**Table S2.** Sex ratio of both sexes embryos injected with cas9 (wildtype) and the sgNb1 (cas9+sg1_ndrg1b)
